# Senescent stroma induces nuclear deformations in cancer cells via the inhibition of RhoA/ROCK/myosin II-based cytoskeletal tension

**DOI:** 10.1101/2021.11.17.468902

**Authors:** Ivie Aifuwa, Jude M. Philip, Byoung Choul Kim, Teresa R. Luperchio, Angela Jimenez, Tania Perestrelo, Sang-Hyuk Lee, Nick Longe, Karen Reddy, Taekjip Ha, Denis Wirtz

## Abstract

The presence of senescent cells within tissues has been functionally linked to malignant transformations. Here, using tension-gauge tethers technology, particle-tracking microrheology, and quantitative microscopy, we demonstrate that senescent associated secretory phenotype (SASP) derived from senescent fibroblasts impose nuclear lobulations and volume shrinkage on malignant cells, which stems from the loss of RhoA/ROCK/myosin II-based cortical tension. This loss in cytoskeletal tension induces decreased cellular contractility, adhesion, and increased mechanical compliance. These SASP-induced morphological changes are in part mediated by lamin A/C. These findings suggest that SASP induces a defective outside-in mechanotransduction, from actomyosin fibers in the cytoplasm to the nuclear lamina, thereby triggering a cascade of biophysical and biomolecular changes in cells that associate with malignant transformations.

## INTRODUCTION

Cellular senescence is a process of cell-cycle arrest that is characterized by a loss of proliferative capacity in mitotic cells ^1–3^. Senescence is triggered in numerous cell types in response to a range of stressors, including replicative, genotoxic or oncogenic stresses. Studies have shown that overexpressing the oncogenes *KRAS,* or the induction of DNA double strand breaks, either by ionizing radiation or other strong mitogenic signals, can induce senescence ^4–8^. Under certain conditions, when damage is irreparable, cells halt their progress through the cell-cycle to limit the perpetuation of cellular damage ^9,10^.

While seemingly beneficial from a tumor-prevention standpoint, senescence can induce critical problems to the overall well-being of organisms. The accumulation of senescent cells within aging tissues result in tissue dysfunctions ^11,12^ that in part stems from soluble factors secreted by senescent cells, termed senescence-associated secretory phenotype (SASP)^7^. In addition, a wide range of transcriptional changes accompany senescence, including significant increases in the secretion of cytokines, chemokines, proteases and growth factors ^7,13,14^. Utilizing *in vivo* and *in vitro* models, SASP has been shown to stimulate survival and proliferation of tumor cells ^15–17^, enable tumor cell invasion through ECM-degrading enzymes such as MMPs ^17^, and promote tumor-driven angiogenesis through the expression of vascular endothelial growth factor (VEGF) ^18^. Furthermore, SASP can drive epithelial-to-mesenchymal transition (EMT) ^7,19^ and enable the dissemination and migration of cells to distal sites. Recently, we demonstrated that SASP promotes cancer cell migration by reorganizing cytoskeletal proteins through the modulation of RhoA activity^20^. Since cytoskeletal proteins are implicated in the shaping of interphase nuclei^21^, we asked whether SASP directly influenced nuclear architecture, mechanics, and associated gene expression.

Alterations and defects in nuclear morphology are associated with numerous human pathologies ^22^, including Hutchison Gilford progeria syndrome ^23^, Werner Syndrome, and more commonly cancer ^12,24^ and aging ^12,25,26^. Among the morphological changes observed in tumor cells, defects in nuclear morphology and architecture remain a key diagnostic feature of malignant cells ^24,27^. These include changes in nuclear size and shape, as well as changes in chromatin organization. The spatial arrangement of the nucleus is maintained by the nuclear matrix, which consists of the nuclear lamina and a network of proteins and RNA ^27^. The most abundant nuclear matrix proteins are lamin A/C and lamin B, which compose the nuclear lamina. The nuclear lamina aids in organizing chromatin within the nucleus, where lamin and lamina-associated proteins bind directly to DNA sequences called lamina-associating domains (LADs) ^28^. Because the nuclear lamina directly interacts with chromatin, changes in nuclear shapes have been shown to alter chromatin organization and gene positioning, thereby influencing gene expression patterns and cellular function ^29^. Nuclear shape is tightly coupled to cytoskeletal elements via linker of nucleoskeleton and cytoskeleton (LINC) complexes and can therefore be influenced by changes in intracellular (inside-out) and extracellular (outside-in) forces ^30^. This occurs through a process called mechanotransduction, where adhesion receptors such as integrins and cadherins are physically coupled to cytoskeletal filaments, linking the nuclear membrane to nucleoli, chromatin and DNA ^31^.

Here, we show that SASP secreted by senescent stromal fibroblasts substantially alter the nuclear morphology of cancer cells. The nuclei of SASP-exposed cancer cells form deep invaginations and exhibit large-scale volume shrinkage. These changes in nuclear shapes, which are also accompanied by changes in underlying molecular patterns, are highly dependent on intact lamin A/C-mediated connections and the reduction in acto-myosin tension. The loss of this cellular tension decreases the lateral pulling forces exerted on the nucleus thereby leading to membrane relaxation and the formation of nuclear lobulations. We demonstrate that RhoA/ROCK-mediated acto-myosin tension is both required and sufficient to induce nuclear-shape changes.

## RESULTS

### SASP induces changes in nuclear morphology and volume of cancer cells

To assess the impact of SASP on the nuclear properties of carcinoma cells, we induced senescence in human WI-38 lung fibroblasts via exposure to bleomycin and collected the SASP-containing media ^20^. This conditioned media (CM) derived from senescent cells was added to human T47D epithelial breast carcinoma cells. Upon exposure, we observed and quantified cellular responses by way of changes in nuclear morphology. Prior to the addition of conditioned media from senescent fibroblasts (denoted as Sen CM), most cancer cells displayed a round morphology with less than 5% of cells displaying an elongated morphology. However, when exposed to Sen CM (48h), approximately 70% of the cells developed an elongated cellular morphology, and 30% retaining a rounded morphology^32^.

To visualize real-time morphological changes in cells exposed to Sen CM, we first utilized phase-contrast microscopy (**Figure 1a**). Most cells switched from a rounded to an elongated morphology, with increased cell extensions and significant decreases in cell-substrate contact area and circularity (**Figure 1b-c** and **supplementary Fig 1a-b**). Furthermore, we observed an increase in the integrated intensity measured from phase contrast micrographs within the traced cell region (**Figure 1a-c**). This apparent optical change indicated that cells underwent significant changes cell height along the z-direction, leading us to ask whether SASP exposure also influenced nuclear morphology.

**Figure 1.**
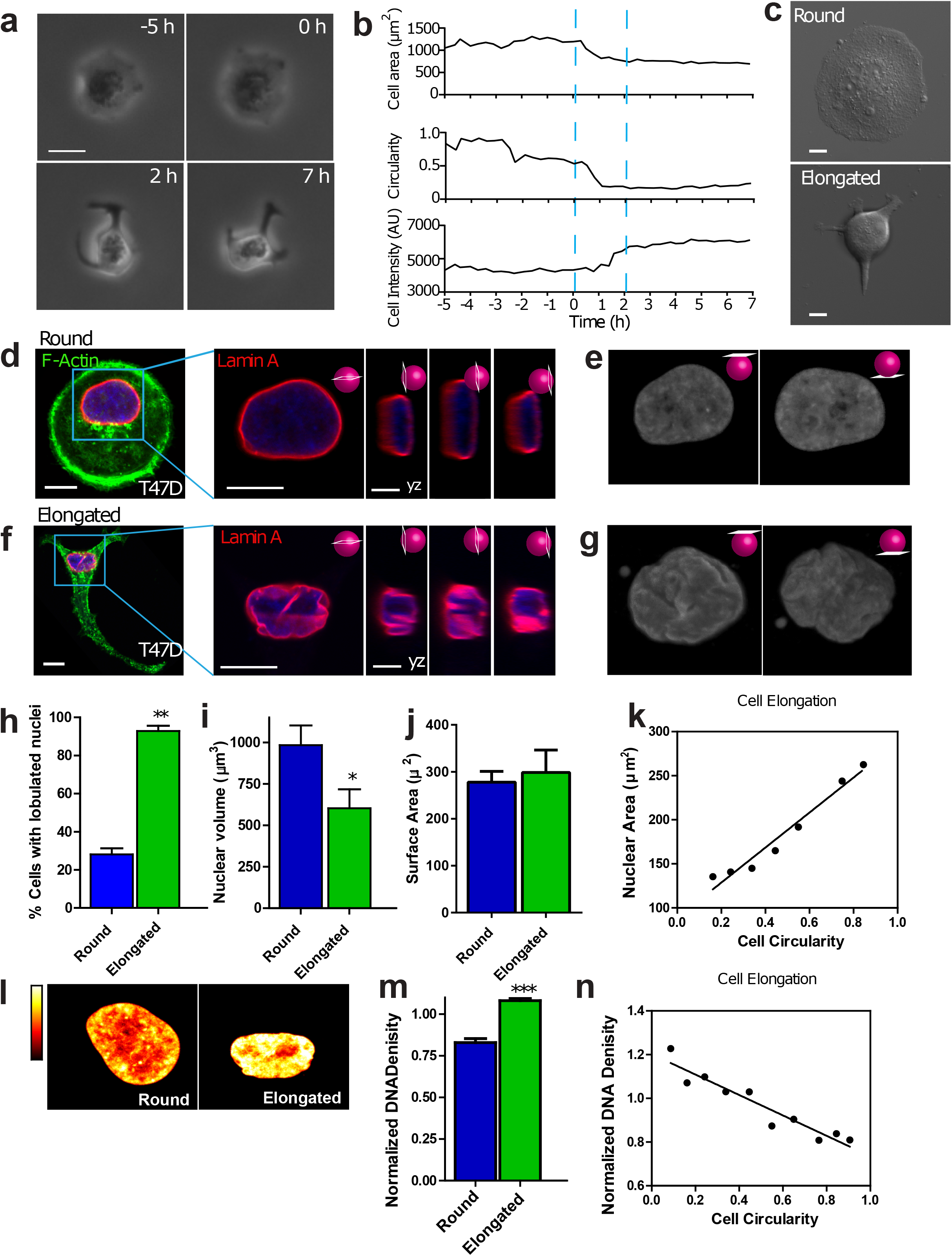
SASP induces changes in cell and nuclear morphology. **(a)** Phase contrast micrographs of a T47D breast cancer cell undergoing a dramatic change in morphology induced by Sen CM. Scale bar, 20 μm. **(b)** Changes in cell area, cell circularity, and cell intensity for the cell shown in (a). (**c**) phase contrast micrographs showing examples of rounded and elongated T47D cells after exposure to Sen CM for 48h **(d** and **f)** Fluorescence confocal micrographs of round (d) and elongated (f) T47D cells stained for F-actin (green), Lamin A (red) and nuclear DNA (blue) after exposure to Sen CM for 48h. Blue squares indicate zoomed regions of T47D cells. Scale bar, 10 μm. Following panels are zoomed in xy and yz cross-sections of the left panel. These include the xy equatorial cross-section, yx left, equatorial, and right cross section respectively. Scale bar, 5 μm. **(e** and **g)** Surface rendering of lamin-A-stained nuclei in panels d and f. Left and right panels indicated top and bottom views of the nuclei, respectively. **(h)** Percentage of cells of round (blue) and elongated cell morphology with lobulated nuclei 48h after exposure to Sen CM. Numbers of examined cells are n=24 and n=43 for round and elongated cells, respectively. **(i)** Nuclear volume of cells having round and elongated cell morphology using consequential z-slices; n=8 and n=10, respectively. **(j)** Nuclear surface area of round and elongated cells; n=8 and n=10, respectively. **(k)** Binned correlation between cellular circularity and nuclear area. Blue triangle indicates a decrease in cell circularity from left to right (elongated to round). **(l)** Intensities of DNA staining were color coded from high to low (white, yellow, orange, red, black). Condensed regions have higher fluorescence intensity compared to the less condensed regions. **(m)** Normalized nuclear density between round and elongated cells. Normalization was performed among repeats (n=87 and n= 157 for round and elongated cells, respectively). **(n)** Correlation between cellular circularity and normalized DNA density. Blue triangle indicates a decrease in cell circularity from left to right (elongated to round).

To better understand this, we collected z-stacks of fluorescent images using confocal microscopy and then reconstructed cells in 3D to visualize rounded versus Sen CM-induced elongated cells (**Figure 1d-g**). Cellular and nuclear morphologies were visualized based on F-actin and nuclear lamin-A staining. In the presence of Sen CM, the nuclei of rounded cells featured a smooth disk-like morphology, whereas the nuclei of elongated cells displayed large deformations, with deep invaginations (**Figure 1e** and **1g**). These nuclear deformations were readily visualized through y-z cross-sections and top/bottom surface views of stained nuclei (right panels in **Figure 1d** and **1f**). In the presence of Sen CM, we observed that ~25% of rounded T47D cells displayed nuclear lobulations, while ~90% of elongated cells displayed nuclear lobulations (**Figure 1h**). The extent of lobulations was determined based on the number of folds present as visualized via lamin A staining, nuclei with ≥2 nuclear folds were deemed lobulated (see methods and supplemental information). Similarly, nuclear deformations were observed in tumorigenic, non-invasive human breast epithelial MCF7 cells exposed to Sen CM (**Supplementary figure 1a-b**).

In conjunction with lobulated nuclear morphology, Sen CM-induced elongated cells displayed a ~50% reduction in their projected nuclear area (**Supplementary figure 1c).** To determine whether this reduction in nuclear area corresponded to a similar reduction in nuclear volume, we utilized a scanning-laser confocal and integrated nuclear areas of serially sectioned, GFP-lamin A-labeled nuclei^33^. We measured a 40% reduction in nuclear volume in elongated cells compared with control round cells (**Figure 1i**). This volume decrease occurred with no significant changes in the surface area of the nucleus (**Figure 1J**), presumably because of the observed extensive folding and wrinkling of the nuclear envelope. Despite this large decrease in nuclear volume, the height of the nuclei of elongated cells significantly increased (**Supplementary figure 1d**), explaining the observed overall bulging observed under phase contrast microscopy, when cells were exposed to Sen CM (**Figure 1a-c**).

As cells transitioned from a rounded to an elongated morphology, they displayed large decreases in cell contact area accompanied by the formation of cell extensions (**Figure 1a-c**). However, because nuclear shape changes seemed to occur synchronously with cellular shape changes, we next asked whether changes in cell morphology correlated with changes in nuclear morphology. We found that the projected nuclear area decreased as cell circularity decreased. i.e. rounder cells featured larger nuclei (**Figure 1k**). Similarly, larger cells featured larger nuclei (**Supplementary figure 1e**). This suggests that SASP-induced nuclear morphological changes may in part be driven by cellular morphology changes.

Large-scale changes in cell morphology have been shown to not only influence nuclear shape, but also affect chromatin remodeling and compaction ^34^. To determine whether SASP-induced changes in intranuclear architecture, we examined the organization of chromatin utilizing DNA staining via Hoechst 33342. This DNA fluorochrome stoichiometrically stains DNA, allowing us to determine cell cycle status, as well as chromatin distribution and condensation ^35,36^. Brightly stained regions indicate increased compaction denoting heterochromatic regions, while dimly stained regions indicate euchromatic regions, denoting less compaction. Significant reorganization of chromatin accompanied nuclear deformation. Nuclei of elongated cells had regions with significantly higher DNA staining compared to control cells (**Figure 1l**). DNA density (total intensity of DAPI-stained DNA divided by nuclear area) was increased ~25% in elongated cells compared to control cells (**Figure 1m**), which increased correlated with progressive elongation in cell morphology (**Figure 1n**).

Together, these results suggest that SASP-induced nuclear deformation and DNA condensation occur in conjunction with changes in cellular morphology.

### Coupled mechanical forces generated by F-actin are required for nuclear shaping

The actin cytoskeletal network is a major mediator of nuclear remodeling processes ^29,37^ in response to changes in cellular morphology ^21^. To determine whether Sen CM elicited changes in filamentous actin structures, we compared cells exposed to fresh medium (FM) and Sen CM. Cells exposed to Sen CM displayed reduced F-actin (**Figure 2a-c**), accompanied by the disappearance of the rich cortical actin structures observed in FM (**Figure 2a-b**).

**Figure 2.**
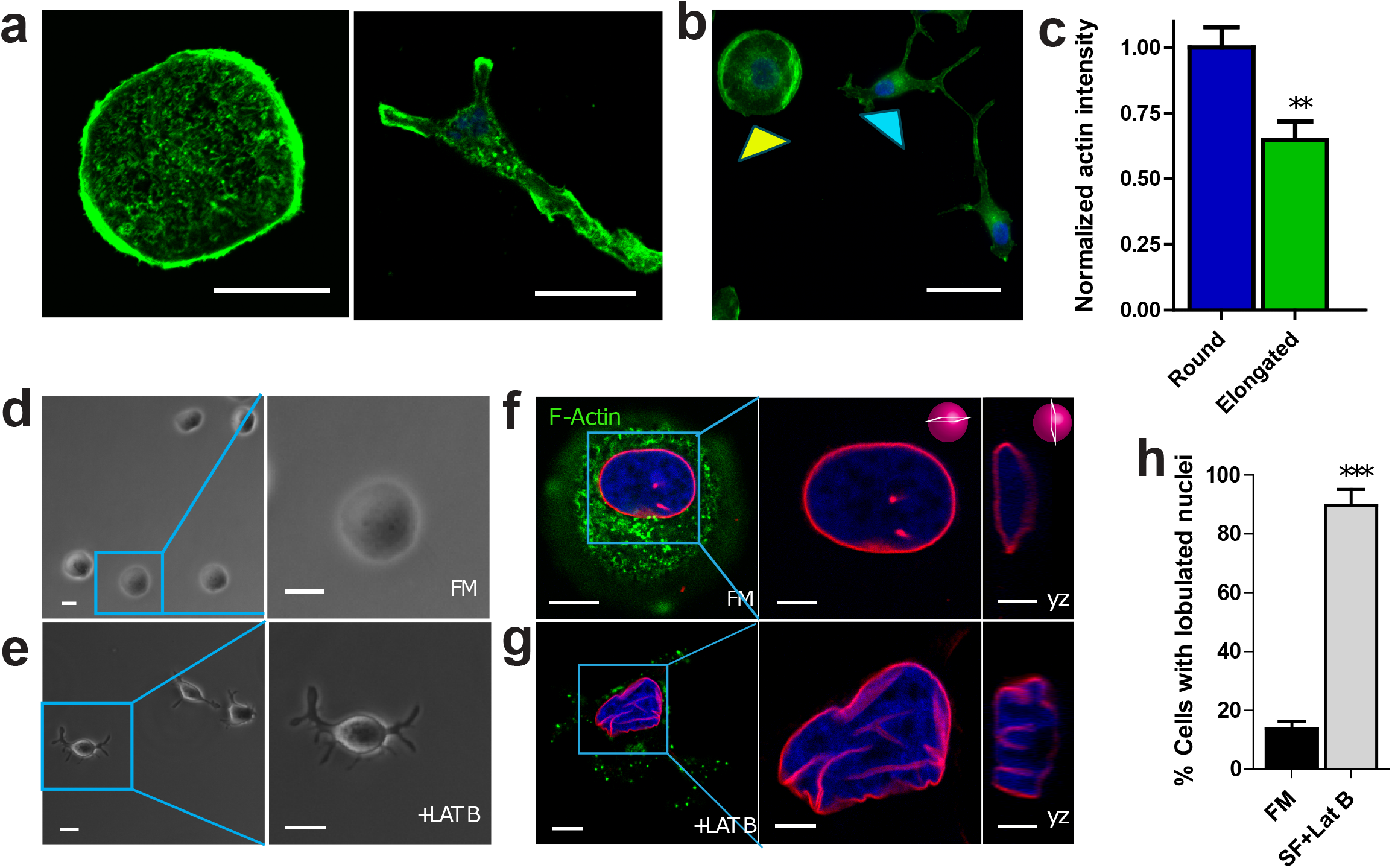
Nuclear shape requires intact F-actin structures. **(a-e)** Fluorescence confocal and phase contrast micrographs of T47D cells after 48h exposure to FM and FM+250 μM of actin-depolymerizing drug Latruculin B, respectively, for 48h. Cells are stained for F-actin (green), Lamin A (red) and nuclear DNA (blue). Scale bar, 10 μm. Panels to the right of f and g show zoomed in xy and yz cross-sections of the left panels. Scale bar 5 μm. **(h)** Percentage of cells featuring a lobulated nucleus 48h after exposure to FM (black) or FM+Lat B (grey). Numbers of examined cells are n=92 and n=49 for FM and FM+Lat B stimulated cells, respectively.

To determine and probe the role of the actin cytoskeleton in regulating normal nuclear morphology, cells were treated with 250 nM of the small-molecule inhibitor of F-actin assembly, Latrunculin B (LatB) in FM. Similar to Sen CM treatment, LatB-treated cells formed long, thick extensions together with significant bulging in the nuclear region (**Figure 2e**) relative to control cells in FM (**Figure 2d**). To visualize the effects of cellular morphological changes on nuclear morphology, cells were stained for F-actin, and Lamin A (**Figure 2f-g**). In addition, LatB-treated cells formed gross nuclear lobulations with a similar proportion of cells having nuclear lobulations as cells exposed to Sen CM (**Figure 2h**).

These findings indicate that LatB-dependent F-actin remodeling elicits similar nuclear morphology changes to Sen CM-treated cells.

### Lamin A/C mediates SASP-induced nuclear changes

While B-type lamins are required for cell viability, A-type lamins (lamin A/C) are major nuclear-scaffolding proteins, which are required for the maintenance of nuclear structure and organization. Lamin A/C plays a pivotal role in mechanotransduction, and the depletion of lamin A/C disrupts the physical connectivity between the cytoplasm and nucleoplasm in normal cells ^38^. LINC complex proteins (linkers of nucleoskeleton and cytoskeleton) are SUN and KASH-domain-containing proteins, which constitute mechanical tethers that span the nuclear envelope and connect a network of lamins and lamin-associated proteins in the nucleus to the cytoskeleton in the cytoplasm ^29,39^. Hence, we asked whether SASP-induced nuclear deformations occurred due to direct connection between the cytoskeleton and nucleoskeleton through lamin A/C.

We found that the expression and localization of Lamin A, nuclear envelope protein Emerin, and LINC complex proteins SUN1, Nesprin2giant, Nesprin3 were largely unchanged following exposure of cells to Sen CM (**Supplementary figure 2a-b)**. These results suggest that Sen CM-induced nuclear invagination is not necessarily due to a loss in expression or mislocalization of these key nuclear-envelope proteins.

Next, we depleted cells of lamin A/C and then exposed to Sen CM (**Supplementary Figure 3a**). We visualized nuclear folds via staining of nuclear-lamina-associated protein Lap2β. Interestingly, lamin A/C-depleted cells exposed to Sen CM failed to alter their nuclear shape (**Figure 3a-d** and **supplementary figure 3b-f**). While 80% of control cells featured nuclear lobulations in response to Sen CM stimulation, less than 5% of lamin A/C-depleted cells formed nuclear lobulations in Sen CM (**Figure 3e**). Although the nuclei of lamin A/C-depleted cells did not form lobulations in FM, an increase in nuclear height was still observed. These results indicate that nuclear lamin proteins - lamin A/C and associated nucleoskeleton-cytoskeleton connections are required for the wrinkling process, *i.e.* without lamin A/C the nucleus fails to wrinkle in response to SASP exposure.

**Figure 3.**
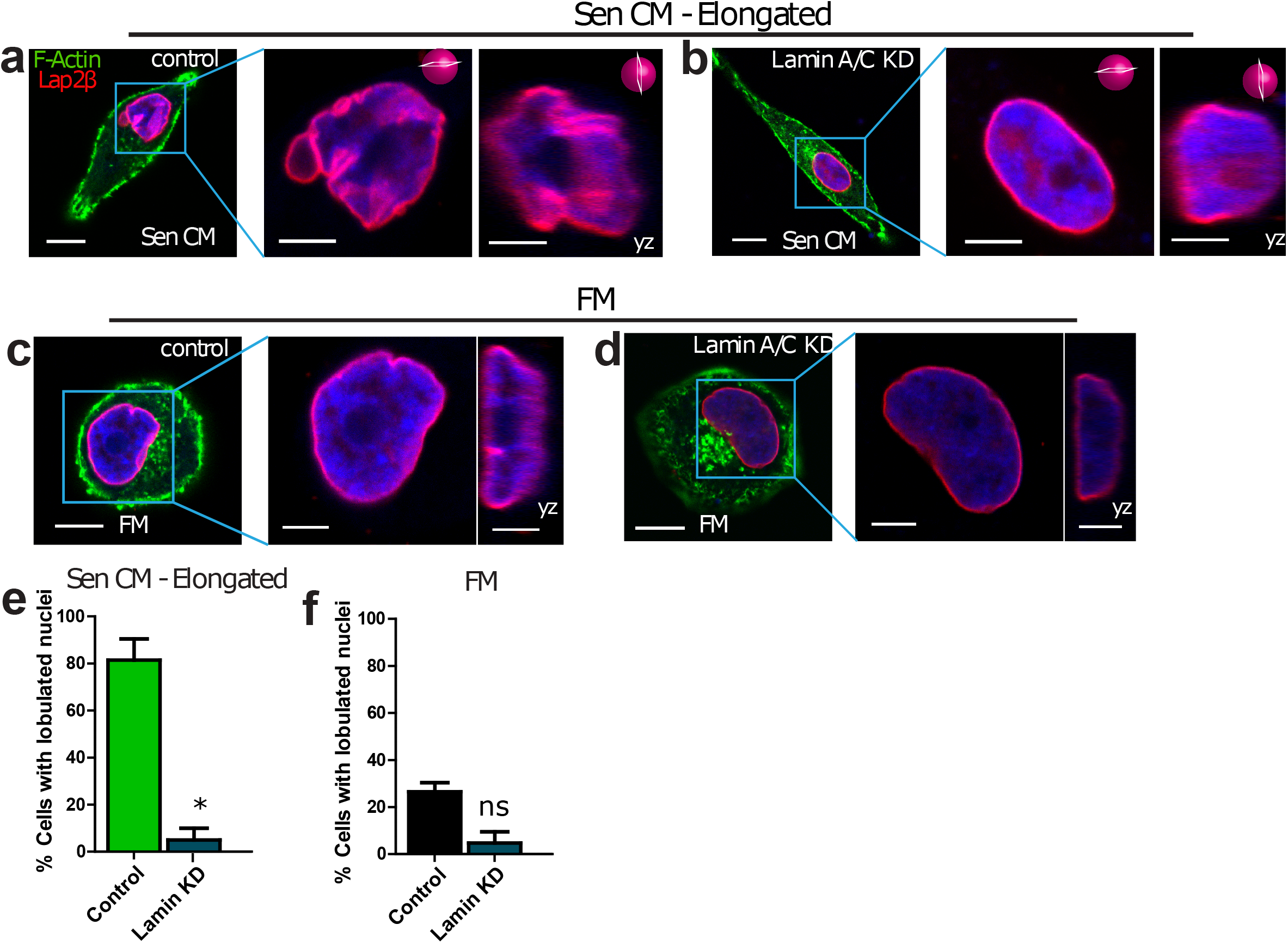
Lamin A/C mediates SASP-induced nuclear changes. **(a-d)** Fluorescence confocal micrographs of control and lamin A/C knockdown cells after stimulation with Sen CM and FM for 48h. Cells were stained with phalloidin, Lap2β, and Hoechst 33342 DNA stain. The following panels are zoomed in xy and yz cross-sections of the left panel. Scale bar 5 μm. These include the xy and yz equatorial cross-sections. **(e-f)** Percentage of control and lamin A/C knockdown cells having lobulated nuclei 48h after exposure to Sen CM (e) and FM (f). n=62, n=33, n=43, n=31 for, Sen CM, Sen CM + lamin KD, FM and FM+lamin KD, respectively.

Because lamin A/C contributes to the maintenance of the structural integrity of the nucleus, we also asked whether lamin A/C deficiency was sufficient to induce nuclear shape changes in FM. We compared the nuclear morphology of control and lamin A/C-depleted cells in FM. In both conditions, nuclei maintained a round morphology (**Figure 3c, 3d, 3f** and **Supplementary figure 3c and f**). This suggests that, although lamin A/C is a major mediator of nuclear shape changes induced by SASP, the depletion of lamin A/C alone is not sufficient to produce SASP-induced changes in nuclear morphology.

### Decreased cytoskeletal tension and cellular contractility drive nuclear shape changes

Cytoplasmic mechanical compliance (deformability) critically depends on nuclear-cytoskeletal connections^40^. To measure cytoplasmic compliance, we utilized particle tracking microrheology ^41^. Carboxylate-modified 100-nm fluorescent polystyrene beads were ballistically injected into the cytoplasm of T47D cells. Since the sizes of the beads are significantly larger than the effective mesh size of cytoskeletal network (20-40nm), the movements of beads reflect the viscoelastic properties of the cells in which they are embedded ^42,43^. Cells were then exposed to Sen CM for 24 h. Using the time-dependent coordinates of the bead-trajectories, we calculated the ensemble-averaged mean square displacement (MSD) of beads embedded within cells exposed to either FM or Sen CM (**Figure 4a-d)**. We observed that particles lodged in the cytoplasm of Sen CM-exposed cells had significantly larger displacements, at all measured time-lags, compared to particles in round cells in FM (**Figure 4e-h**). Together, these results indicate that elongated cells exposed to Sen CM are more deformable than cells exposed to FM, which is consistent with reduced F-actin content in cells exposed to Sen CM (**Figure 2a-c**).

**Figure 4.**
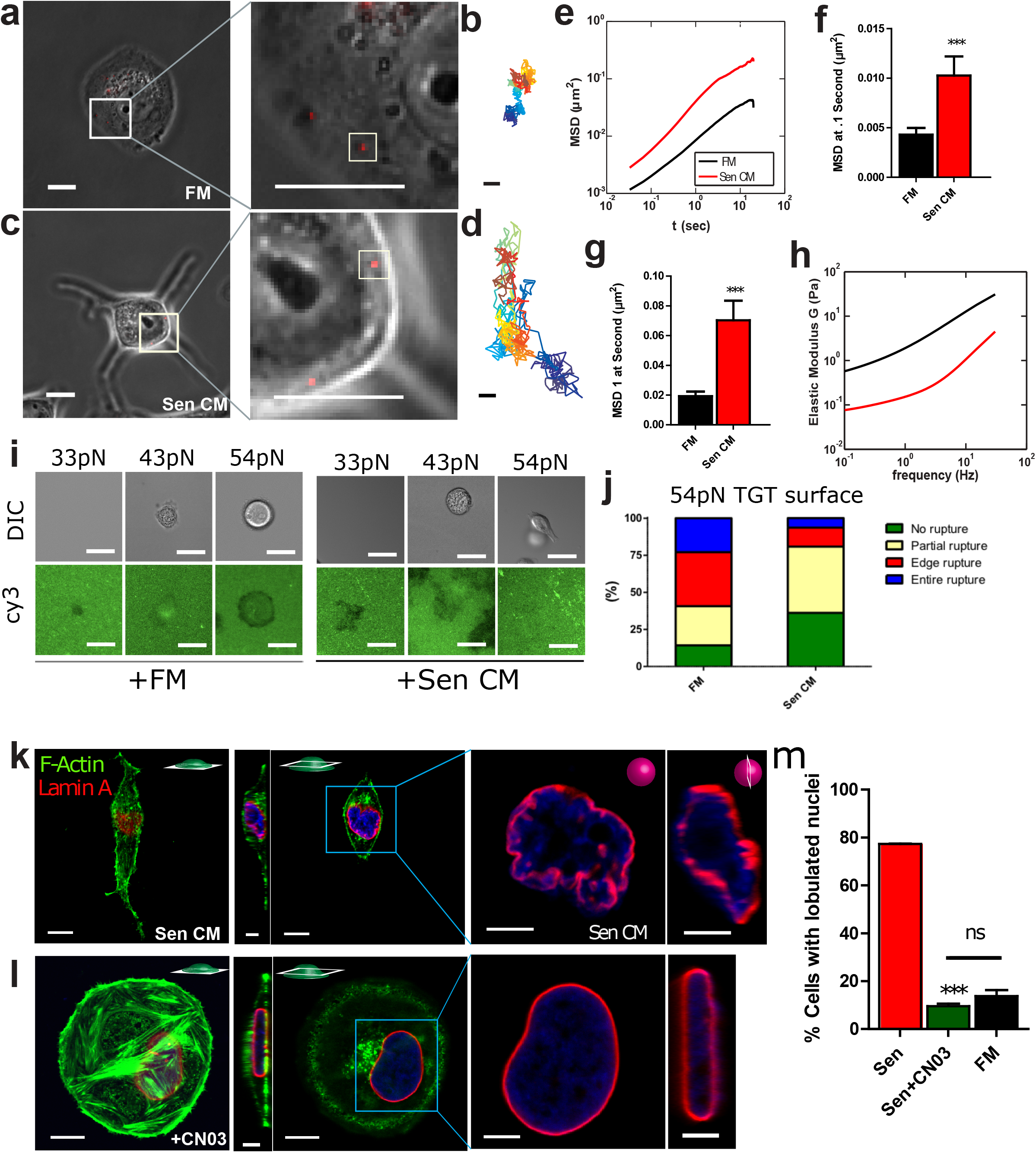
SASP-induced nuclear shape is accompanied with decreased cell contractility and increased cell deformability. **(a, c)** Phase contrast micrographs of T47D cells bombarded with 100nm-diameter fluorescent beads. Post bombardment, cells were exposed to FM (a) and Sen CM (c) for 48h. Scale bar, 20 μm. White squares indicate the regions of each cell expanded to better display the embedded fluorescent particles in the following panel. Second panel shows a zoomed view of 100 nm-diameter fluorescent beads within the cytoplasmic region of the cell in the previous figure. White squares indicate individual particles within each T47D cell. Scale bar, 5 μm. **(b, d)** Particle trajectories of beads indicated in the second panel of (a) where passage of time is indicated by color change. Beads were tracked with ~10 nm spatial resolution and 33 ms temporal resolution for 20 s. **(e)** Population-averaged mean squared displacement (MSD) of particles within the cytoplasm of round and elongated T47D cells as a function of time lag. Numbers of examined cells are n=85 and n=48, for FM and Sen CM conditions, respectively. (**f-g**) Population-averaged MSD evaluated at time lags of 0.1 s (f) and 1 s (g). **(h)** Computed Elastic modulus of cell exposed to FM or Sen CM. (**i**) FM and Sen CM-exposed T47D cell adhesion on TGT surfaces with different tension tolerances (33, 43 and 54 pN) after 3 hr incubation. Images show adherent T47D cells on TGT surfaces and TGT rupture patterns. A loss of Cy3 signal indicates the molecular force larger than tension tolerance of each TGT was exerted by cells. (**j**) Percentage of rupture patterns of T47D cells exposed to FM or Sen CM. n = 92 for FM, n = 65 for Sen CM. Entire rupture indicates TGT rupture in the entire cell projected area. Edge rupture indicates TGT rupture at the cell periphery. Partial rupture indicates any other small ruptures underneath a cell. **(k)** Fluorescence confocal micrographs of T47D cells after 48h exposure to Sen CM and Sen CM+ 1.0 μg/ml Rho activator II CNO3. Cells were stained for F-actin (green), Lamin A (red) and nuclear DNA (blue). First and third panels are basel and equatorial xy cross section of imaged cells. Scale bar, 10 μm. Second panels display yz cross sections. Scale bar, 5 μm. Blue squares indicate zoomed regions of T47D cells in the following panels. The last panels are xy and yz equatorial cross sections of nuclei in previous panels. Scale bar, 5 μm. **(i)** Percentage of cells with lobulated nuclei 48h after exposure to Sen CM (red; n=53 tested cells), Sen CM+CN03 (green; n= 54), and FM (black; n= 92).

Cells exposed to Sen CM display decreased number and size of focal adhesions (**Supplementary Figure 4a-b**) and decreased traction forces on the underlying matrix compared to control cells ^20^. For control cells, maximal traction forces and adhesions are formed at the cellular cortex. Because traction forces exerted by cells are governed by a cell-intrinsic surface balance of contractile forces ^44^, we sought to determine whether elongated cells were less contractile than their round counterparts ^45,46^. To assess cellular cortical tension at molecular levels, we employed the tension gauge tethers (TGT) assay^47^ where a ligand for cell membrane receptors (RGD for integrins in this case) is tethered to a surface through a rupturable DNA tether. Polymer-passivated glass substrates coated with RGD-conjugated TGTs having tension tolerance *T*_tol_ of 33, 43 or 54pN were used. For *T*_tol_ = 33 pN, under exposure to both FM and Sen CM, cells did not adhere stably, and imaging of fluorescence Cy3 labels on the tether showed fluorescence loss, likely because the cells failed to experience > 40 pN forces needed to activate adhesion response through single integrin-ligand bonds due to TGT rupture ^47^. For *T*_tol_ = 43 pN, both FM and Sen CM-exposed cells adhered stably on the TGT surface as expected^47^ and retained a round morphology. For *T*_tol_ = 54 pN, cells exposed to FM remained round with significant TGT rupture in cell periphery or cell center) (**Figure 4i-j and Supplementary figure 4d**), indicating that cells can apply tension stronger than 54 pN, likely through large focal adhesions^48^. In contrast, cells exposed to Sen CM developed an elongated cell morphology with minor ruptures mostly at the front and back extensions, supporting a reduction in cortical cell tension for cells exposed to Sen CM relative to FM.

Next, we asked whether increased contractility would prevent nuclear-shape changes. In non-muscle cells, contractility is modulated via myosin II, primarily by phosphorylation of myosin regulatory light chain ^49,50^. Myosin light chain (MLC) phosphorylation can occur directly via a myriad of proteins including myosin light chain kinase (MLCK) and Rho-associated coiled-coiled containing kinase (ROCK), an effector of RhoA GTPase ^51^. Therefore stimulation of the Rho-Rock pathway induces myosin II-mediated contractility and tension ^52^. Hence, to increase cell tension and contractility, we increased RhoA activity by treating cells exposed to Sen CM with CN03 (constitutive activator of RhoA). In the presence of Sen CM and in response to increased RhoA activity, cells retained their rounded morphology, preserved prominent actin stress fibers and a high number of large focal adhesions, and prevented SASP-induced nuclear deformations (**Figure 4k-m**, **Supplementary figure 4c**). These results suggest that the reduction in RhoA-based contractility mediates SASP-induced nuclear lobulations.

### Reduction in cell contractility is sufficient to induce nuclear deformation and gene expression changes

We asked whether the reduction in cellular tension and contractility alone was sufficient to recapitulate SASP-induced nuclear deformation. We treated cells with inhibitors of various proteins of the acto-myosin network, including CT04 (FM+CT04), Y27632 (FM+Y27632) and blebbistatin (FM+Bleb) to inhibit RhoA, ROCK, and myosin II ATPase activity in FM, respectively. Treated cells not only acquired an elongated morphology ^53^, but also developed nuclear deformations and invaginations (**Figure 5a**). Cells with inhibited RhoA, ROCK, and myosin II activity had similar fractions of cells with lobulated nuclei as cells exposed to Sen CM (**Figure 5b**). Since myosin II is a direct regulator of cellular tension, we asked whether blebbistatin treatment could recapitulate the phenotypic effects of Sen CM. In line with previous reports that cells with pronounced myosin contractility typically harbor large focal adhesion complexes ^50^, we observed decreased staining of focal adhesion protein vinculin after blebbistatin treatment (**Supplemental Fig 5a-b**). Furthermore, we found that blebbistatin-treated cells displayed a similar decrease in cell spreading and nuclear area (**Supplemental Fig 5c-d**) as Sen CM stimulated cells ^20^.

**Figure 5.**
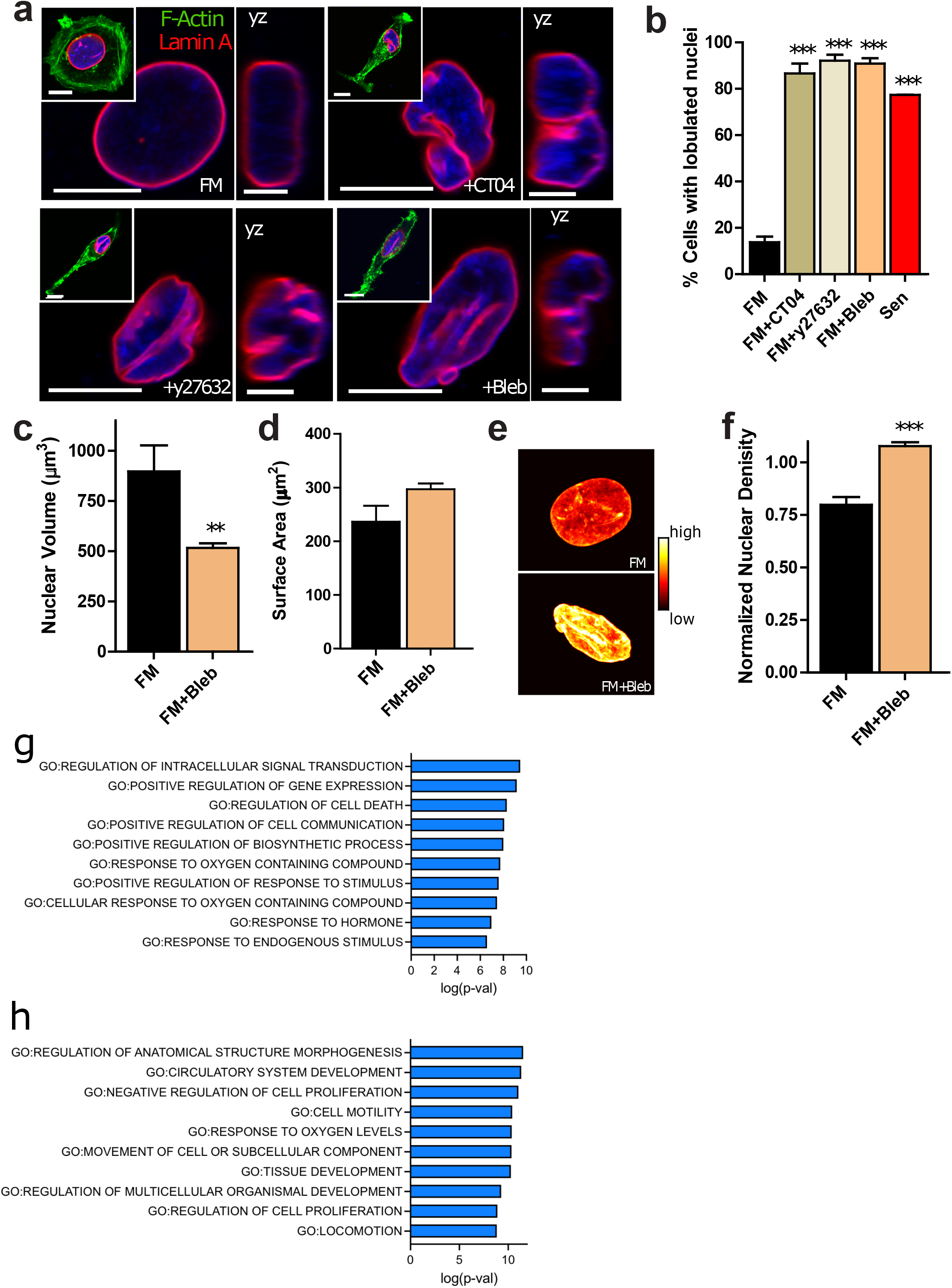
Reduction in cell contractility alone is sufficient to induce nuclear deformation following SASP stimulation. Fluorescence confocal micrographs of T47D cells after 48h exposure to FM, FM+2.0 μg/ml Rho Inhibitor I CT04, FM+15 μM ROCK inhibitor Y27632, and FM+25 μM myosin II inhibitor Blebbistatin (Bleb). Cells are stained for F-actin (green), Lamin A (red) and nuclear DNA (blue). Images display xy and yz equatorial cross-sections respectively. Scale bar, 5 μm. Whole cell mid-plane images of are displayed as insets. Scale bar, 10 μm. **(b)** Percentage of single cells with lobulations when exposed to Sen CM (red; n=53), FM (black; n=92), FM+CT04 olive; n=74), FM+Y27632 (beige; n=38) and FM+Bleb (orange; n=49) after 48h. **(c)** Nuclear volume of T47D cells after 48h exposure to FM and FM+ 25 μM Blebbistatin - using consequential z-slices; n=13 and n=15 respectively. **(d)** Nuclear surface area of FM and FM+ Blebbistatin stimulated cells; n=13 and n=15 respectively. **(e)** Intensities of DNA staining per condition were color coded from high to low (white, yellow, orange, red, black). Condensed regions have higher fluorescence intensity compared to the less condensed regions. **(f)** Normalized nuclear density between FM and FM+ Blebbistatin stimulated cells. Normalization was performed among repeats (n=43 and n= 112 for FM and FM+ Blebbistatin stimulated cells, respectively). (**g-h**) Bar plots showing the transcriptional programs for overlapping upregulated (g) and downregulated (h) genes between Sen CM and FM+Bleb treated cells.

Next, measuring nuclear volume and nuclear surface area, we observed that myosin II inhibition induced a similar reduction in nuclear volume without a significant change in nuclear surface area, with a compensating increase in nuclear height, indicating similar nuclear wrinkling and bulging phenotypes (**Figure 5c-d and Supplementary Fig 5e**) to Sen CM exposed cells. Furthermore, measuring chromatin compaction after myosin II inhibition, we observed the characteristic increase in nuclear density via H33342 staining (**Figure 5e-f**). Together these results suggest that the inhibition of intracellular tension and contractility are necessary (**Figure 4**) and sufficient (**Figure 5**) to recapitulate the nuclear phenotypes induced by Sen CM.

Given the significant deformations exhibited by Sen CM exposed nuclei, we asked whether cells harbored similar gene expression patterns when treated with myosin II inhibitor—blebbistatin. Using bulk RNA sequencing of cells treated with FM, Sen CM, and FM+Bleb, we identified 1726 genes for Sen CM and 471 genes for FM+Bleb which changed significantly from FM condition (**Supplementary Figure 6a-d**). Comparing the overlapping downregulated and upregulated genes in both Sen CM and FM+Bleb treated cells, we found 168 and 71 genes, respectively. Using Cytoscape and Cluego, we found that the upregulated genes involved in the Sen CM condition corresponded to programs involving the cellular responses to stress and lipids, and cell death; whereas the downregulated genes associated to pathways involving regulation of cell adhesion and cell-cell communication (**Supplementary Figure 6e-f**).

Next, we determined the transcriptional programs shared among the upregulated and downregulated genes in both the Sen CM and FM+Bleb conditions. We rationalized that these overlapping genes likely provide insight into the genes associated with the induced nuclear morphological changes in both conditions. Using gene set enrichment analysis (GSEA), we asked what cellular programs were linked to the overlapping upregulated and downregulated genes. We found that the upregulated genes were involved in processes involving intracellular signal transduction, regulation of gene expression and cell death, and cell communication (**Figure 5g**). Downregulated genes involved processes linked to cellular structural morphology, proliferation, migration and locomotion (**Figure 5h**).

Together these results suggest that changes in gene expression induced by Sen CM exposure are partially shared with cells exposed to FM+Bleb, and that changes in the nuclear morphology, intracellular compliance and cytoskeleton-based contractility have profound effects on the gene expression patterns in malignant cells.

## DISCUSSION

SASP produced by senescent fibroblasts in the stromal space and nearby connective tissues of tumors can alter the cellular phenotypes of neighboring cancer cells. These phenotypes include enhanced invasion and proliferation, as well as EMT ^7,15,16,54,55^. Recently, we have demonstrated that SASP stimulation facilitates the reorganization of cytoskeletal proteins, which induces onset of migration in natively non-motile cancer cells ^20^. Here, we observe that the nuclei of Sen CM-stimulated cells display large-scale lobulations and deep nuclear invaginations. The nuclei of Sen CM-stimulated cells also display a large decrease in volume accompanied by an increased height with unchanged surface area^33^. This indicates a wrinkled nuclear phenotype with contraction in the lateral direction but an expansion in the normal direction, thereby suggesting a loss of lateral and apical mechanical support. Along with nuclear shape changes, SASP stimulation promotes chromatin reorganization and condensation. These SASP-stimulated alterations in nuclear shape and chromatin compaction correlate well with global cell morphology changes, indicating that these events occur concurrently.

SASP-mediated nuclear lobulations and invagination - and associated decreased in nuclear volume - critically depend on F-actin assembly and acto-myosin contractility. Numerous studies have demonstrated that cell shape information is transduced into nuclear shape changes via mechanical forces transmitted by the actin cytoskeleton ^21,37,56^. External forces distort specific adhesion molecules such as integrins, which link to actin microfilaments through focal adhesions ^57^. In fibroblasts and several other types of normal cells, nuclear shape modulation via actin is mediated in large part by a dome-like contractile acto-myosin structure, called the perinuclear actin cap ^21,30,38^. Dorsal actin fibers associate with LINC complex proteins at the apical surface and sides of the nucleus and promote dynamic anchoring between the cytoskeleton and the nuclear interior through lamin A/C ^21,58,59^. However, apical cross sections of both round and elongated cancer cells displayed punctate actin structures on the top and sides of nuclei, with no apparent nucleus-associated fiber structures resembling the perinuclear actin cap. However, nuclear shape is also regulated by central actin filaments pulling laterally on the nuclear membrane via anchor proteins and terminating with adhesion proteins ^37,45^. To assess the role of actin cytoskeleton, we first determined the role of nucleus-cytoskeletal connection in regulating SASP-stimulated nuclear deformation, we depleted cells of lamin A/C and assessed the response. While Sen CM exposure leads to a reorganization of the cytoskeleton in both control and lamin-depleted cells, lamin A/C-depleted cancer cells in FM failed to modulate their nuclear shape, unlike normal cells. This suggests that lamin A/C and intact nucleus-cytoskeletal connections are required for the formation of nuclear invaginations in cancer cells in response to SASP stimulation. This also suggests that cell shapes and cytoskeletal forces are transmitted into nuclear deformations via lamin A/C, further strengthening the paradigm that transmitted extracellular forces influence nuclear morphology and chromatin organization.

While intact nucleus-cytoskeletal connections are required for the transmission of forces from the cytoskeleton to the nuclear interior, nuclear shape changes occur through the fluctuations and changes in intracellular and extracellular forces ^29,56,60^. Because extracellular forces remained unchanged between FM and Sen CM stimulated conditions, we measured the intracellular mechanics of the cells. Utilizing particle-tracking microrheology, we determined that Sen CM stimulated cells were considerably more deformable than control cells. We also observed reduced number and size of focal adhesions ^20^ and a significant reduction in traction forces in Sen CM ^20^. Together these results indicate that Sen SC stimulation facilitates the loss in cellular tension and contractility. Therefore, we speculated that this decreased actomyosin force-profiles acting on the nucleus may result in the nuclear wrinkling and formation of invaginations. To evaluate this, we stimulated cellular contractility/tension by the constitutive activation of upstream regulator of myosin II, Rho A. With RhoA stimulation, nuclei remained round and disk-shaped in the presence of Sen CM. This result suggests that the loss of tension/contractility is required for SASP induced nuclear deformation.

We asked whether a loss in cell tension/contractility was sufficient to promote a nuclear phenotype similar to SASP stimulation. To evaluate this, we inhibited Rho/ROCK/myosin II activity and assessed nuclear morphology. Inhibition of cell tension was sufficient to recapitulate nuclear phenotypes similar to SASP stimulation, where nuclei developed a similar wrinkling and chromatin condensation. Together, these suggests that the decrease in cell tension/contractility are both necessary and sufficient to induce SASP-induced nuclear deformation.

Lastly, we aimed to understand the role of Sen CM in regulating genome function and to determine the contributions of cell and nuclear morphological change on this process. Although Sen CM stimulation influences cell and nuclear morphology, additional factors in the conditioned medium may independently influence gene expression. To observe the contribution of cell and nuclear morphology on gene expression, we assessed the expression profile with blebbistatin treatment and compared the changes to that of Sen CM. We observed significant changes in the transcription programs of cells exposed to Sen CM and FM+Bleb, with 161 overlapping downregulated genes and 73 overlapping upregulated genes relative to FM controls. GSEA analysis of the overlapping genes indicated that there were significant downregulation of cellular processes involving regulation of cellular morphogenesis, cell motility and cellular proliferation, indicating that SASP stimulation is biochemically and biophysically sufficient to alter gene expression.

To explain these findings, we propose a model by which actomyosin tension exerted on the nucleus mediated by lamin A/C together with alterations in nuclear-cytoplasmic force balance creates an outside-in physical signaling cascade. With control cells (FM), the actomyosin forces pull laterally on the nucleus and flattens it into a disk shape (**Supplementary Figure 6g**). Here, actomyosin forces are balanced by mature focal adhesions enabling a rounded cellular morphology. However, SASP stimulation reduces RhoA activity ^20^. This in turn reduced myosin II-mediated contractility, as observed by the reduction of cell traction forces^20^ and cell adhesion. Because, RhoA mediates the maturation and turnover of focal adhesions through myosin II ^61^, the reduction of cytoskeletal tension and reduction in cellular adhesions promote a decrease in the actomyosin forces laterally pulling on the nucleus. Moreover, reduction in cytoskeletal tension also alters the nucleoplasm-cytoplasm mass transport and pressure difference, which results in the loss of water and protein content in the nucleus and the formation of folds and invaginations within the nucleus walls^33^. This reduction in nuclear mass helps to propagate the observed alterations in cellular and nuclear mechanics. These factors together trigger a cascade that leads to the force-dependent changes in projected gene expression profiles.

## ACKNOWLEDGEMENTS

This work was supported in part by the National Institutes of Health Grants U54CA143868 (DW), U01AG060903 (DW, JMP), National Science Foundation grant PHY1430124 (to T.H.), and by the Johns Hopkins University Older Americans Independence Center of the National Institute on Aging (NIA) under award number P30AG021334 (JMP). T.H. is an investigator of the Howard Hughes Medical Institute.

## AUTHOR CONTRIBUTIONS

I.A. and D.W. conceived and designed the study. IA, BCK, NL, SHL, TL, AJ and TP performed the experiments. IA, JMP, BCK, TL, NL, and SHL analyzed the results. IA, JMP and DW interpreted results; DW, JMP, KR, and TH supervised the study. IA, JMP, DW, BCK, TL, TH, wrote and edited the manuscript.

## MATERIALS AND METHODS

### Cell culture

T47D and MCF 7 cells (ATCC) were cultured in RPMI 1640 medium (ATCC) and DMEM respectively supplemented with 10% (v/v) fetal bovine serum (Hyclone Standard) and 1% (v/v) Penicillin/ streptomycin (Sigma). Wi-38 (ATCC) cell were cultured in Eagle’s Minimal Essential Medium (EMEM; ATCC) supplemented with 10% (v/v) fetal bovine serum (Hyclone Standard) and 1% (v/v) penicillin/streptomycin (Sigma). Human fibroblasts were rendered senescent as indicated in Aifuwa et al ^20^. Cells were verified to be senescent via Ki-67 immunostaining and a count of population doublings over a period of 4 days^20^. All cells were maintained at 37°C and 5% CO_2_ in a humidified environment during culture and imaging. T47D cells were seeded unto tissue culture dishes and allowed to adhere and spread for a 24h. Cells were then washed 3x with PBS and incubated with EMEM based FM or CM.

### Conditioned medium

Conditioned medium (CM) from human fibroblasts was prepared by washing cells 3x with PBS and incubating cells with serum-free EMEM for a period of 24 h. Condition medium from these senescent cells (denoted Sen CM in the text) contained IL6 and IL8 indicated in Aifuwa et al ^20^.

### Drug treatments

The microtubule destabilizer nocodazole (Sigma), the F-actin disassembly drug latrunculin B (Sigma), the non-muscle myosin II inhibitor blebbistatin (Sigma), the ROCK inhibitor Y27632 (Sigma), the Rho inhibitor CT04 (Cytoskeleton Inc.), and the Rho activator CN03 (Cytoskeleton Inc.) were diluted from stock using culture medium. Nocodazole was used at a final concentration of 0.5 μg/ml; latrunculin B was used at a final concentration of 250 nM; blebbistatin was used at a final concentration of 25 mM; Y27632 was used at a final concentration of 15 mM; CT04 was used at a final concentration of 2 μg/ml; CN03 was used at a final concentration of 1 μg/ml.

### Immunofluorescence

To visualize Lamin A, microtubules and F-actin, cells were fixed in 4% paraformaldehyde (Electronic Microscopy Systems) for 10 min and permeablized with 0.1%Triton X-100 (Fisher Chemicals) for 10 min. Cells were blocked at room temperature for 1 h in 1% BSA (Sigma). Subsequently, cells were incubated in primary antibodies for 1 h at room temperature. Antibodies used included: Rb. Anti-Lamin A (1:50; ab16667 Abcam), sMs. Anti-Vinculin (1:400; Sigma). Cells were then incubated with either Alexa 568 conjugated donkey anti-mouse or anti-rabbit secondary antibodies, Hoechst 33342 and phalloidin for 1h at room temperature, after which cells were washed with PBS. Fluorescent images were collected after 48 h exposure to FM or CM conditions. Cells were imaged using a Nikon A1 confocal microscope using a 63X oil-immersion lens.

### Tension gauge tether (TGT) assay

The TGT assay was utilized to estimate molecular tension on a single integrin-ligand bond during cell adhesion and spreading of T47D as described previously^47^. In brief, a cyclic peptide RGDfK (cRGDfK) ligand and a Cy3 dye were conjugated to each end of an 18-nucleotide single stranded DNA (5-/Cy3/GGC CCG CAG CGA CCA CCC/cRGDfK/-3). The ssDNA was hybridized with a biotin-tagged complementary ssDNA (5-/GGG TGG TCG CTG CGG GCC/-3) at distinct locations, resulting in a double stranded DNA with different tension tolerances (T_tol_ = 33, 43, and 54 pN). Then, 1 μM of each TGTs was immobilized on a PEGylated glass slide via a neutravidin-biotin interaction. Next, T47D cells at the density of 10^5^ cells/ml were seeded on the TGT surfaces with either the fresh medium (FM) or the conditioned medium from the senescent cells (Sen CM). After 3hr incubation at 37°C, the cells were fixed with 4% para-formaldehyde. The fixed cells and TGT rupture patterns induced by the cells were imaged by an epi-fluorescence microscope (Nikon Ti-E, Nikon Inc.). The number of adherent cells on each TGT surface and the rupture patterns were further analyzed using ImageJ.

### Nuclear volume

Nuclear volume for T47D cells under the various conditions was measured from randomly selected cells stained with Hoechst 33342. Confocal microscopy was used to image nuclei by 0.3 μm z-stacks. Nuclear area per each cell was traced using NIS-Elements image analysis software (Nikon) and nuclear volume was then calculated by summing the nuclear area per 0.3 μm z-stack.

### Image analysis

Image analysis was performed using a customized MATLAB program to segment cell and nuclear boundaries using phalloidin actin stain and Hoechst 33342 DNA stain, respectively. Fluorescence intensity was quantified per pixel within the segmented region. Nuclear density was calculated by summing the total DNA intensity within the traced nuclear region divided by the nuclear are. Cellular circularity was calculated as 4πA/P2. Where A is the area and P is the perimeter of the measured cell.

### Microrheology

T47D cells were plated overnight and allowed to adhere. Particle-tracking measurements are detailed in Wu et al ^41^.

### Gene expression analysis using Cytoscape

RNA sequencing was conducted for both conditions and the fold change versus FM was computed for all assessed genes. Using a significance cutoff for −log(p-value) equal to 3 and the log_2_foldchange greater than 0.5, we determined the significantly upregulated and downregulated genes. Using the list of significant genes, using Cytoscape we inputted the genes into the ClueGO app and using the *Homo Sapien* marker list, GO Biological processes pathway analysis, of pathways </= 0.05 that includes GO Term fusion genes we generated the network maps of the interacting pathways based on the gene lists for both the upregulated and downregulated genes. For the GSEA analysis the overlapping genes were pasted into the online GSEA analyzer and again using GO Biological processes we generated the significantly changed pathways based on the gene lists.

### Statistical analysis

The number of cells and biological repeats (n) for all experiments are indicated in the figure captions. Mean values, standard error and all statistical analyses were calculated and plotted using Graphpad Prism 9 (Graphpad Software, San Diego, CA). One-way ANOVA and unpaired t-tests were conducted to determine significance, depending on the number of variables assessed. Statistical significance (p) was indicated within the figures using the following scale: *** for p < 0.001, ** for p < 0.01, and * for p < 0.05.

## SUPPLEMENTARY FIGURES

**Supplementary Fig 1.**

**(a-b)** Fluorescent confocal micrographs of round (a) and elongated (b) MCF7 cells stained for F-actin (green), Lamin A (red) and nuclear DNA (blue) after exposure to Sen CM for 48h. Left and right panels display xy and yz equatorial cross-sections respectively. Whole cell images are displayed as insets per image. Scale bar, 10 μm. **(c)** Averaged size of cells exposed to Sen CM after 48h. **(d)** Nuclear height of cells having round and elongated cell morphology using consequential z-slices; n=8 and n=10 respectively. **(e)** Binned correlation between cellular area and nuclear area. Blue triangle indicates an increase in cell area from left to right (elongated to round).

**Supplementary Figure 2**

Localization of nuclear lamina protein lamin A, nuclear envelope protein emerin, and LINC complex components Sun2, Nesprin2-giant, and Nesprin 3 in round and elongated T47D cells. Scale bar, 10 μm **(b)** Western blot of T47D cells exposed to FM and Sen CM. The bands correspond to lamin A, Lamin C, Nesprin 2, and β-actin from top to bottom.

**Supplementary Figure 3**

**(a)** Immunoblots of scramble and lamin A/C knockdown T47D cells. Top bands are lamin A and lamin C from top to bottom. **(b-c)** Fluorescence confocal micrographs of control and lamin A/C knockdown (262765) cells after stimulation with Sen CM (b) and FM (c) for 48h. Cells were stained with phalliodin, Lap2β, and Hoechst 33342 DNA stain. The following panels are zoomed in xy and yz cross-sections of the left panel. Scale bar 5 μm. These include the xy and yz equatorial cross-sections respectively. **(d-f)** Percentage of control and lamin A/C knockdown cells having lobulated nuclei 48h after exposure to Sen CM and FM.

**Supplementary Figure 4**

**(a-c)** Fluorescent confocal micrographs of T47D cells exposed to FM (a), Sen CM (b) and Sen CM +CN03 (c) then stained for vinculin, where blue boxes indicate zoomed region of T47D cells. Scale bar, 10 μm in original image and 5 μm in zoomed image. (**d**) distribution plot of cell circularity in each condition, 1 indicates a perfect circle on TGT surfaces.

**Supplementary Figure 5**

**(a-b)** Fluorescent confocal micrographs of T47D cells exposed to FM (**a**) and FM +Bleb (**b**) then stained for vinculin, where blue boxes indicate zoomed region of T47D cells. Scale bar, 10 μm in original image and 5 μm in zoomed image. **(c)** Averaged nuclear area of cells exposed to FM and FM+Bleb after 48h. **(d)** Nuclear height of cells having round and elongated cell morphology using consequential z-slices; n=13 and n=15 respectively.

**Supplementary Figure 6.** Shared changes in gene expression for SASP stimulation and myosin II inhibition. **(a,b)** Volcano plots of log_2_ (fold change) vs. −log(p-value) showing the significantly upregulated and downregulated genes in Sen CM (a) and FM+Bleb treated cells (b). **(c,d)** Venn diagrams showing the number of significant genes unique and shared between Sen CM and FM+Bleb treated cells for both the downregulated (c) and upregulated genes (d). **(e, f)** Network representation of pathways associated with the upregulated (e) and downregulated (f) genes for Sen CM treated cells, generated using Cluego analysis in Cytoscape. (**g**) model highlighting the key role of the actin cytoskeleton in the formation of nuclear invaginations as a result of Sen CM exposure.

