## Supplementary material for "Senescent stroma induces nuclear deformations in cancer cells via the inhibition of RhoA/ROCK/myosin II-based cytoskeletal tension": Supplamentary documents

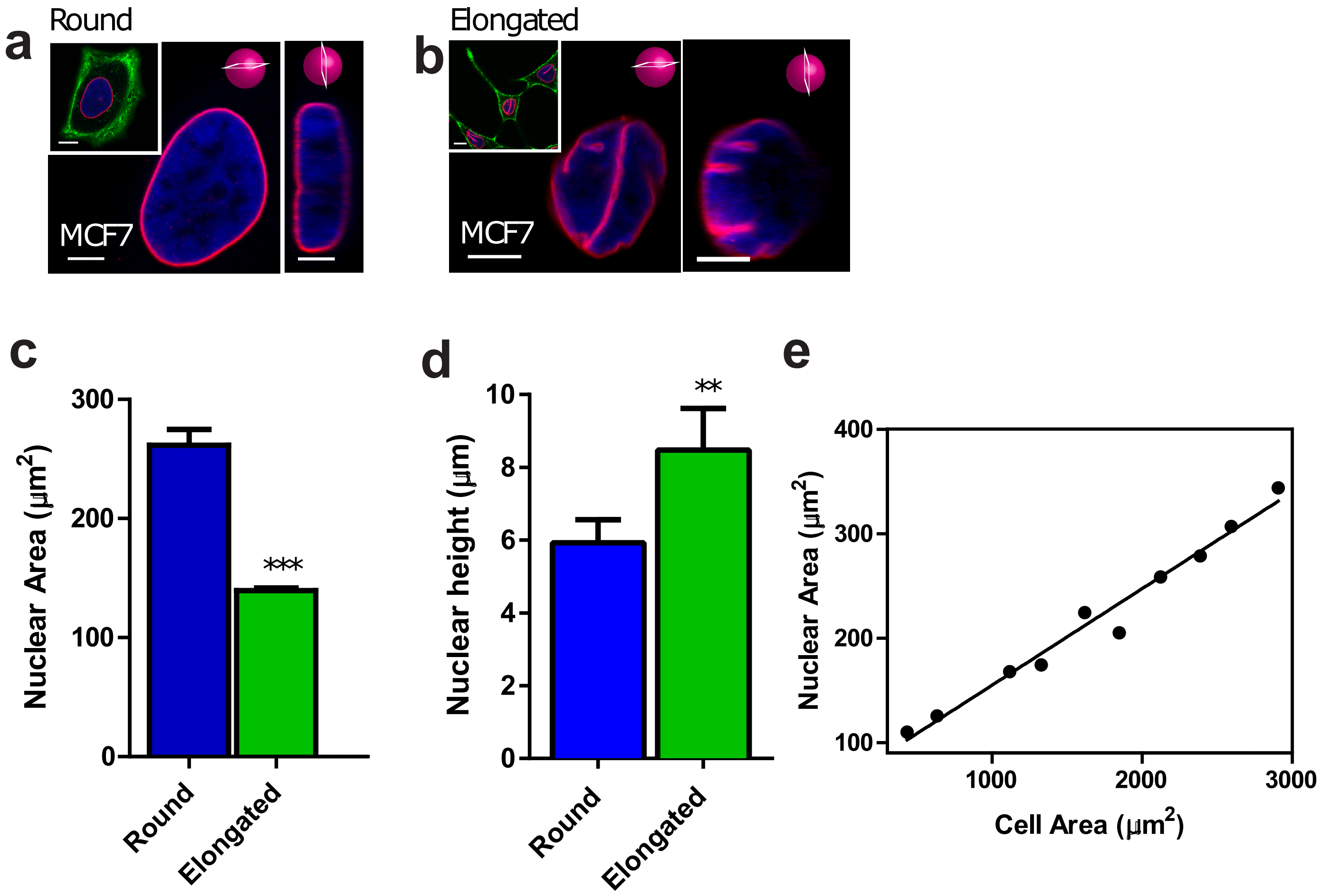


**Supplementary Figure 1**

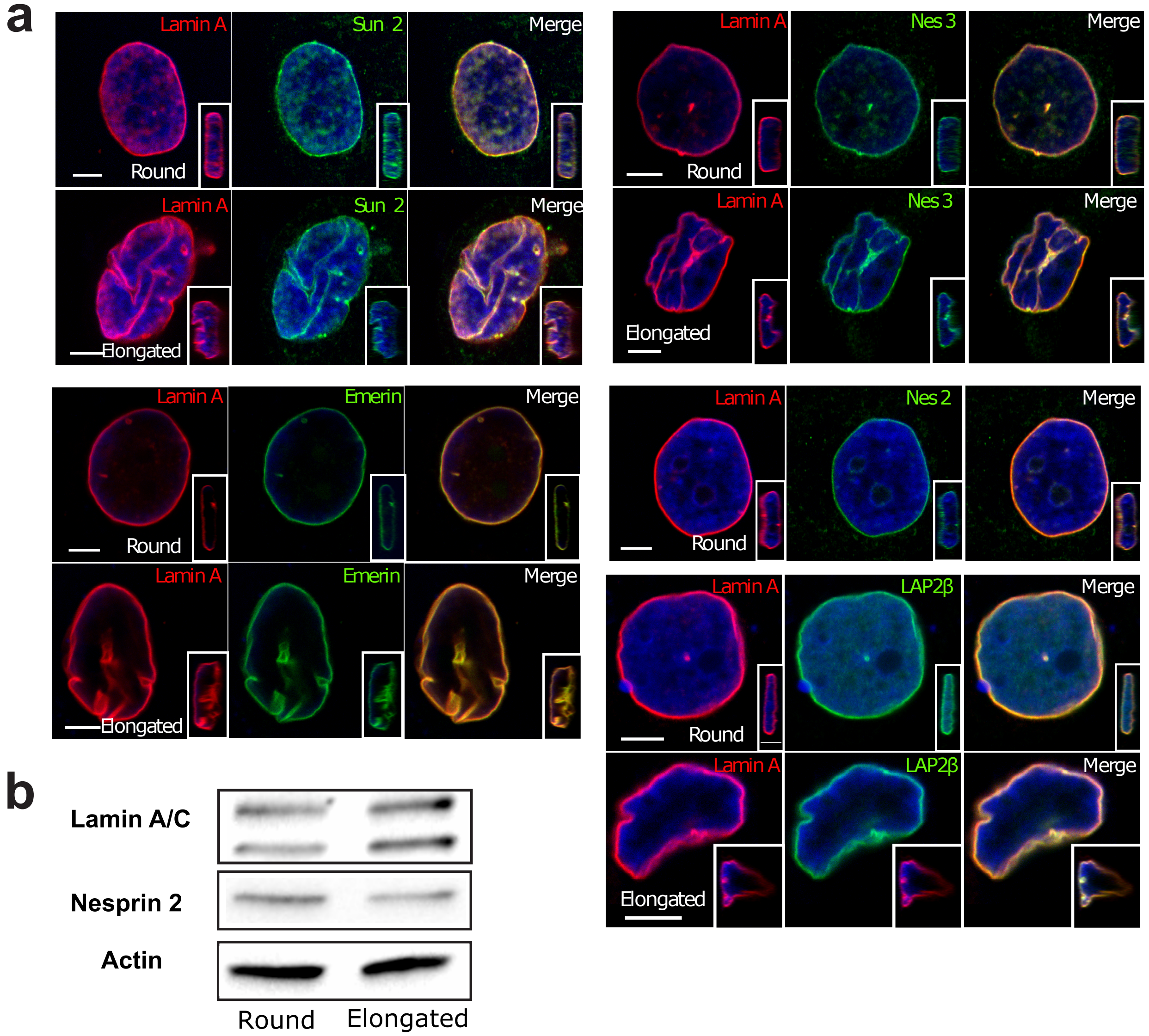


Supplementary Figure 2

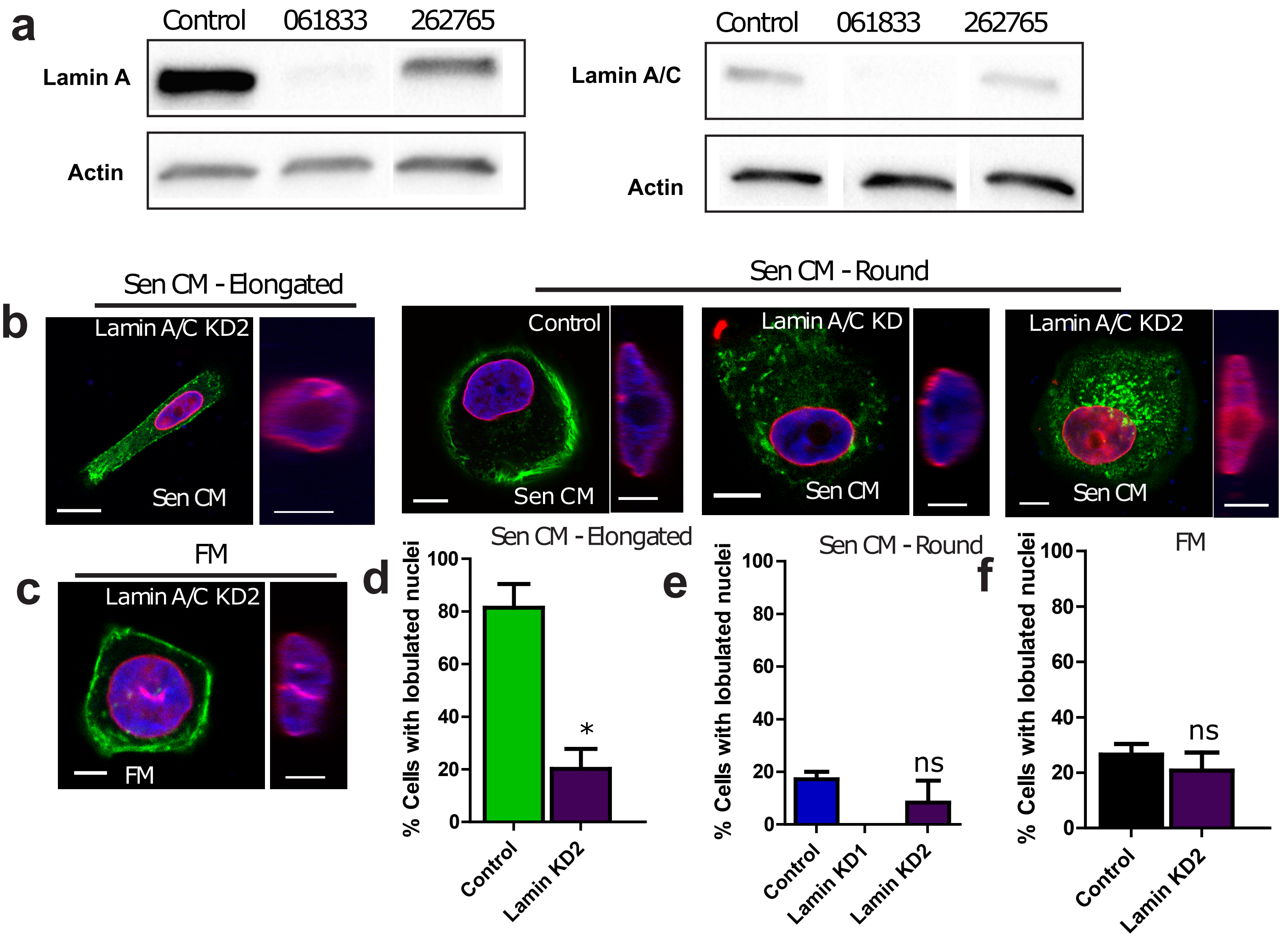


Supplementary Figure 3

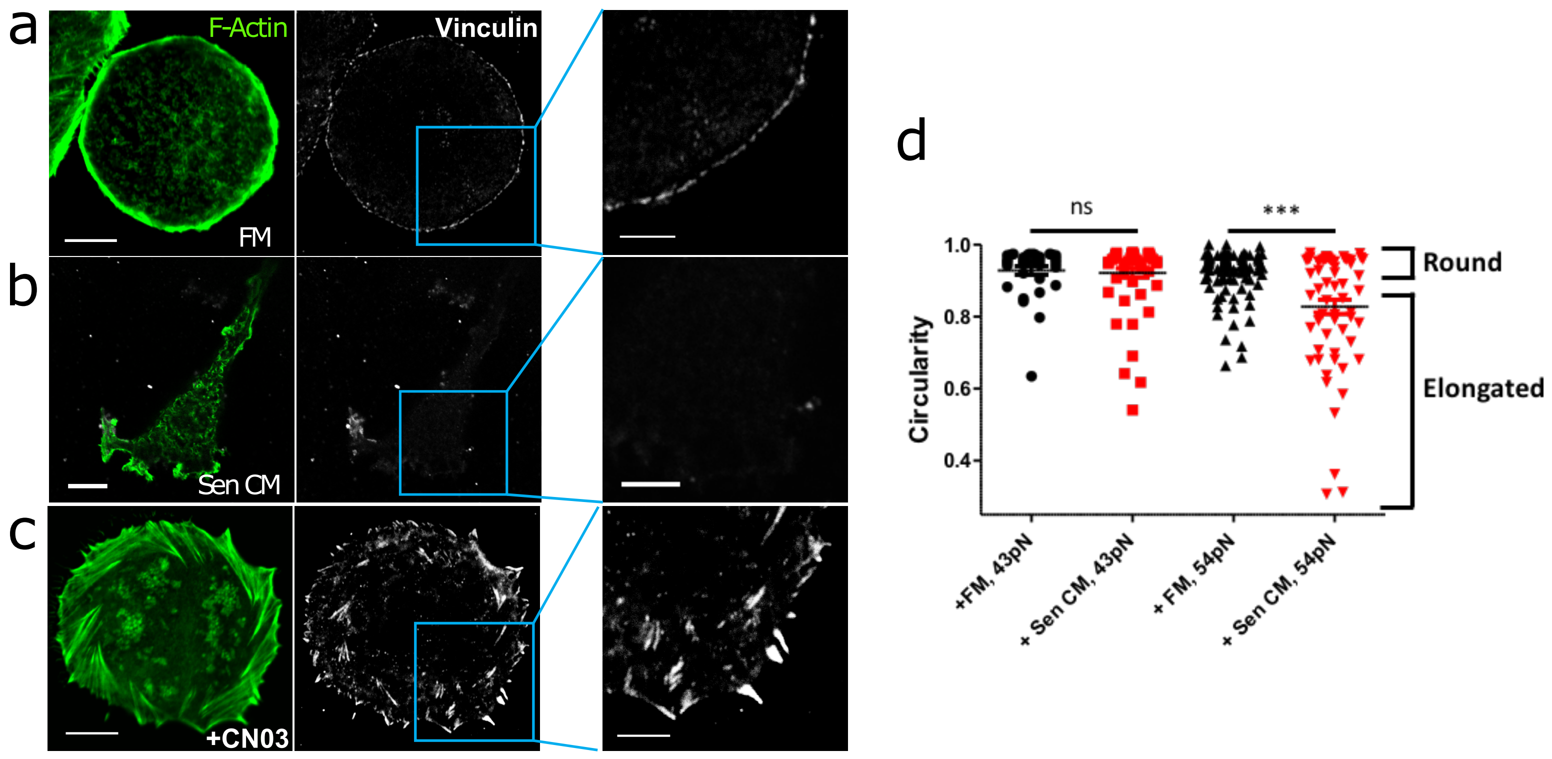


Supplementary Figure 4

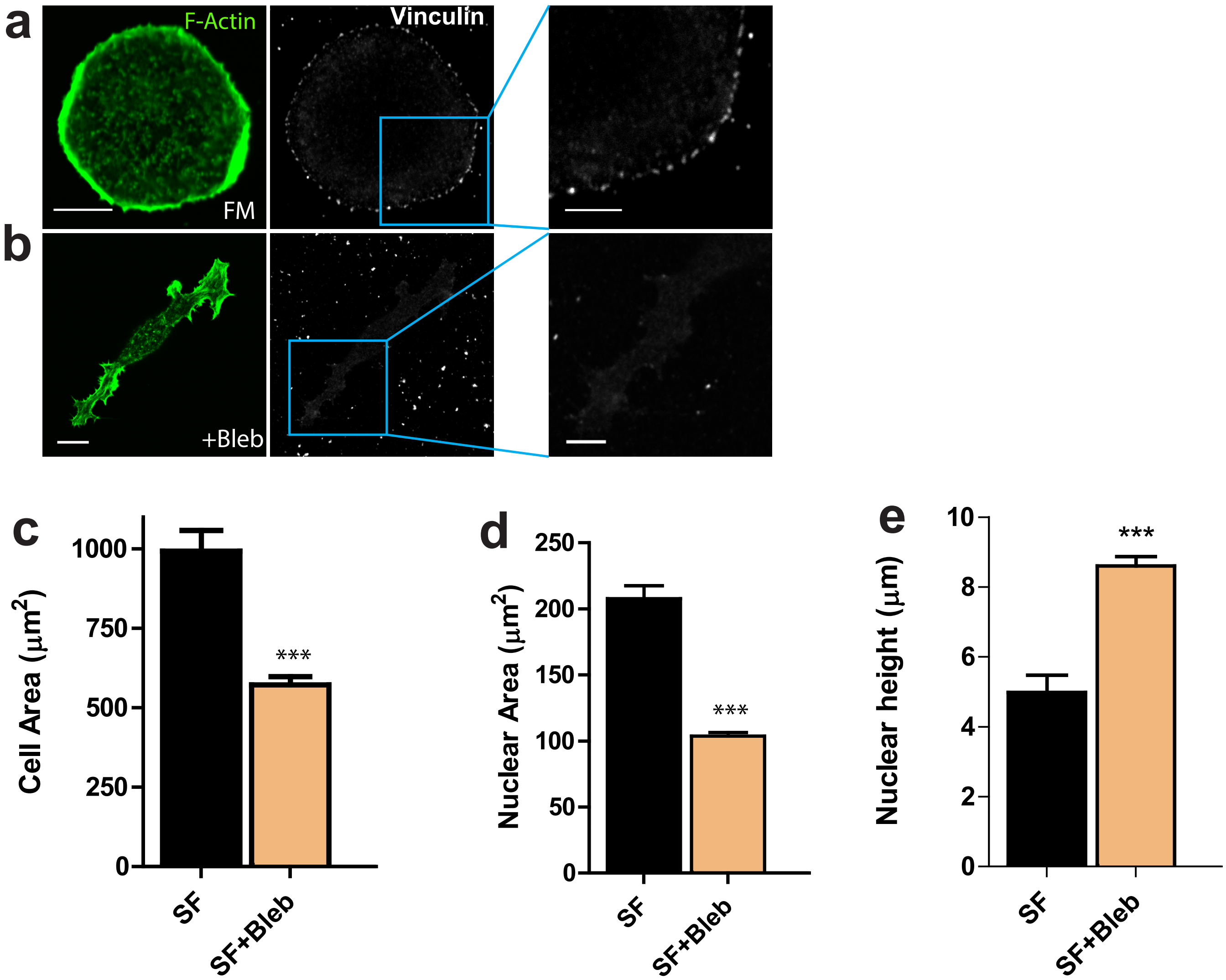
a
F-Actin
Vinculin
FM
b
+Bleb
c
Cell Area (μm²)
0
250
500
750
1000
***
SF
SF+Bleb
d
Nuclear Area (μm²)
0
50
100
150
200
250
***
SF
SF+Bleb
e
Nuclear height (μm)
0
2
4
6
8
10
***
SF
SF+Bleb

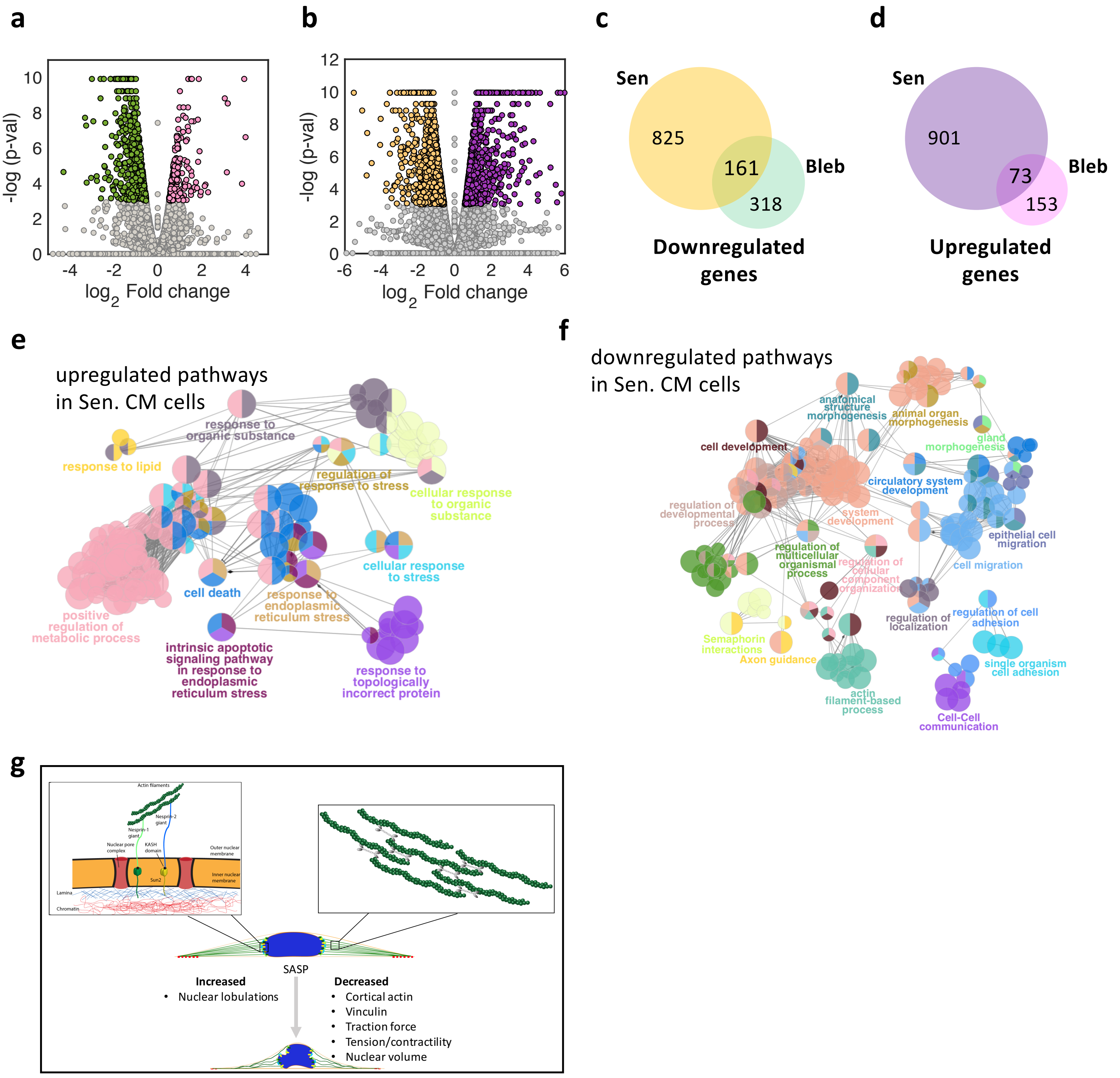


Supplementary Figure 6
